# Scalable production of recombinant three-finger proteins: from inclusion bodies to high quality molecular probes

**DOI:** 10.1101/2022.04.11.487962

**Authors:** Jiang Xu, Xiao Lei, Ao Li, Jun Li, Shuxing Li, Lin Chen

## Abstract

We introduce a working pipeline for expression, purification and validation of disulfide-bond rich three-finger proteins using *E. coli* as the expression host. With this pipeline, we have successfully obtained highly purified and bioactive recombinant α-Bungarotoxin, k-Bungarotoxin, Hannalgesin, Mambalgin-1, α-Cobratoxin, MTα, Slurp1, Pate B etc. Milligrams to hundreds of milligrams of recombinant three finger proteins were obtained within weeks in the lab. The recombinant proteins showed specificity in binding assay and six of them were crystallized and structurally validated using X-ray diffraction protein crystallography. As many three finger proteins have attractive therapeutic or research interests and due to the extremely high quality of the recombinant three finger proteins we obtained, our method provides a competitive alternative to either their native counterparts or chemically synthetic ones, and should facilitate related research and applications.

## Introduction

The snake venom is a large repertoire of digestive enzymes, toxin peptides and compounds. The three-finger neurotoxins (TFNs) are a collection of such toxin peptides used by the snake to kill the prey through binding and blocking ion channels in the neurological system. The α-neurotoxin binds to and block the muscle type nicotinic acetylcholine receptor (α1_2_β1γ(ε)δ nAChR) on the neuromuscular junction, leading to paralysis of the prey. The snake venom also contains various toxin peptides that have interesting properties. Some of them have analgesic effect, such as mambalgin-1 (Diochot et al., 2012) and Hannalgesin (Pu et al., 1995b), some of them can bind to receptors in the neurological system, such as MTα and κ-Bungarotoxin, which bind to Muscarinic α2B-adrenoceptor (Servent and Fruchart-Gaillard, 2009, Näreoja et al., 2011) and α3β2 nAChR (Chiappinelli, 1983; Dewan et al., 1994), respectively. Although these toxin peptides may have attractive usage in biomedical research, not all toxin peptides are present in large quantities in snake venoms. κ-Bungarotoxin (κBtx), for example, only takes a very small fraction of the venom, was often contaminated by α-Bungarotoxin (αBtx), leading to inconsistent results in some of the earlier researches(Chiappinelli et al., 1989), and is commercially unavailable now, thus further restricted their study and usage.

Three-finger proteins (TFPs) are a class of proteins (peptides) including TFNs and their mammalian homologue that show similar three-finger structure as TFNs. These mammalian TFP homologues are referred to as the Ly6/uPAR family proteins, some famous players of which include Lynx (Faure et al., 2016; George et al., 2017; Lyukmanova et al., 2017, 2016; Thomsen et al., 2016), Slurp(Akbar et al., 2019; Fischer et al., 2001; Kryukova et al., 2019; Paramonov et al., 2020; Shah et al., 2016; Swamynathan and Swamynathan, 2016; Taylor et al., 2016), which are proposed to be endogenous modulators of nAChR, the Pate (Prostate and Testis Expression) family (Levitin et al., 2008; Margalit et al., 2012; Rajesh and Yenugu, 2017; Turunen et al., 2011), which play important roles in the capacitation of the sperm and fertilization and CD59, a GPI-anchored membrane protein that protects the cell from complement attack(Huang et al., 2006; Kieffer et al., 1994; Michielsen et al., 2017; Nevo et al., 2013). As the detailed biological function of many of these toxin-like proteins (peptides) still largely remains elusive, it is thus desirable to have a reliable production method for the scientific community.

Due to the complex intramolecular disulfide bonds system in TFPs, it is usually impossible to obtain correctly folded TFPs directly from *E. coli* and the expressed recombinant proteins are always in the form of inclusion body (I.B.), and all previous attempts included an additional refolding process (Antil-Delbeke et al., 2000; Fiordalisi et al., 1991; Rosenthal et al., 1994). Other expression systems were also attempted, such as *Pichia Patoris(Fiordalisi et al., 1996; Krajewski et al., 2001; Levandoski et al., 2000), or Eukaryotic* expression systems(Trémeau et al., 1995). While these attempts obtained recombinant three finger proteins (rTFP) and met the end claimed in the research, all these methods suffered from sophisticated post-purification cleavage of fusion tags and low yield. Until now, very few of these recombinant three-finger proteins (rTFPs) have been structurally validated by X-ray diffraction (XRD) protein crystallography studies. There were successful attempts using chemical synthesis, such as muscarinic toxin MT7 and MT1 (Fruchart-Gaillard et al., 2012), and the pain-killing toxin Mambalgin (Diochot et al., 2012; Pan et al., 2014; Schroeder et al., 2014), but this method suffered from sophisticated synthesizing steps and high costs. As such, a universal, high yield production protocol capable of producing high quality rTFPs is desired.

We previously reported the production of bioactive recombinant α-Bungarotoxin (rec-αBtx) (Xu et al., 2015), in which we used radioimmunoassay to determine the optimal refolding conditions. For other TFPs, without a valid activity determination method, it is usually hard to tell which refolding condition is optimal for a particular TFP. Here we introduce an efficient, productive pipeline that we used to generate various high quality, rTFPs, most of which were structurally and biochemically validated.

## Materials and Methods

The overall experimental design is illustrated in **Figure 1**

**Figure 1.**
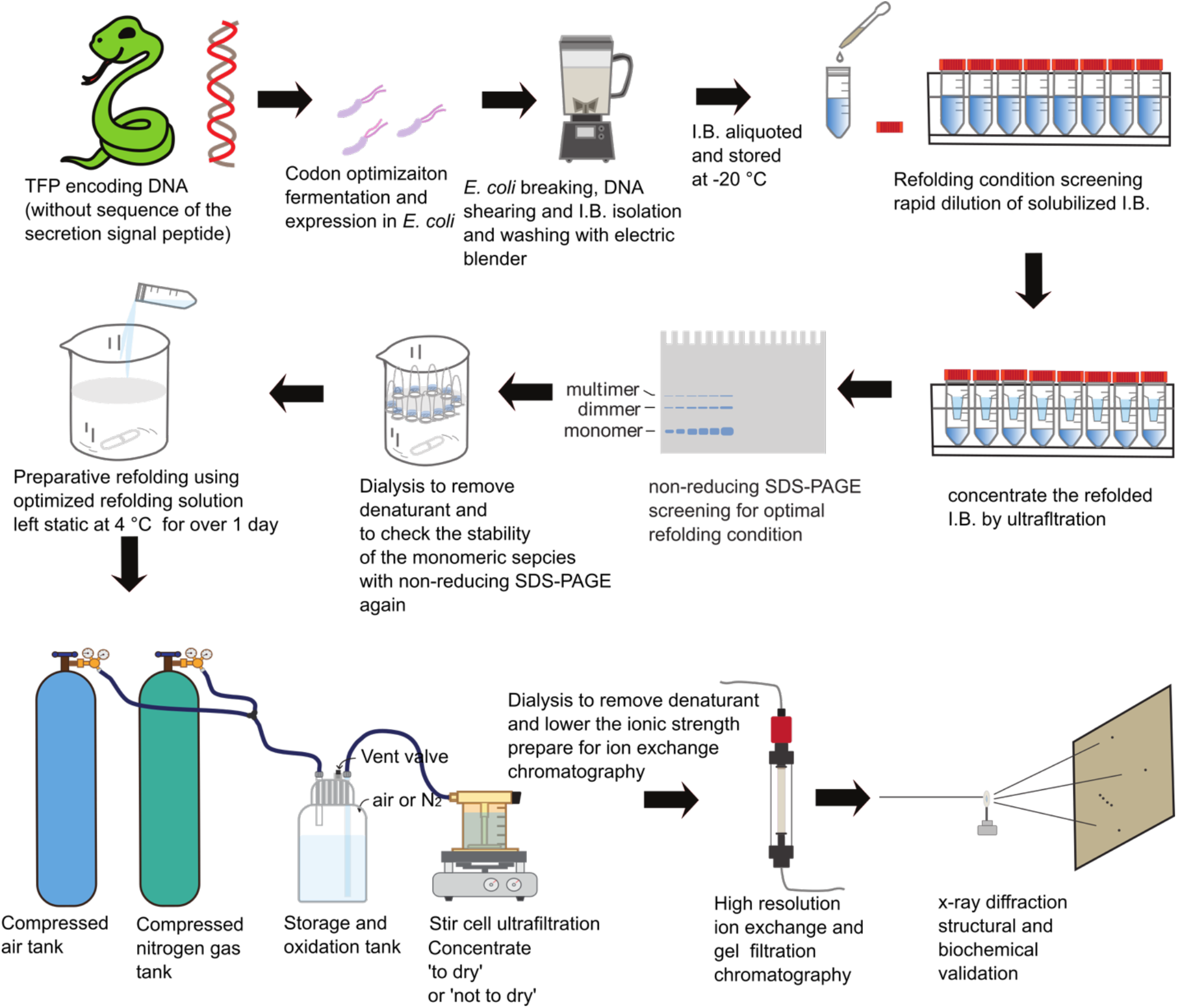
Experimental flow chart illustration.

### Buffers, enzymes, and chemicals

Lysis buffer: 50 mM Tris-HCl(pH 8.0), 150 mM NaCl, 1% Triton X-100, 10 mM 2-mercaptoethanol (2-ME) 5 liter for 200 g of bacteria cell pellets, should be freshly prepared)

Solubilization buffer: 50 mM Tris-HCl (pH 9.0), 6 M guanidine-HCl (or 8 M urea), and 5 mM 2-ME (should be freshly prepared).

Refolding screening buffer (Supplementary table 1)

Restriction enzymes were from Takara or New England Biolabs. All chemicals were from Sigma-Aldrich unless otherwise stated.

### Vector construction, *E. coli* fermentation and inclusion body extraction

Genes encoding the toxin proteins were codon-optimized for expression in *E*. coli and synthesized (Genscript Inc, Integrated DNA Technologies Inc) with NdeI site on the 5’ end and a termination codon (TAA or TAG) at the 3’ end just before the XhoI sites. The genes were inserted into the NdeI and XhoI sites of pET30b (Novagen) and the reconstructed expression vector was transformed into the expression host BL21(DE3). *E. coli* cells were fermented either with a home-made 5-liter fermenter or a BioFlo3000

Bioreactor (New Brunswick Scientific) and induced for protein expression by adding 0.8 mM IPTG at an Optical density of 18∼19 and fermented for additional 4 hrs. Typically, 160∼400 g of bacteria pellets (wet weight) could be obtained and stored at -20 °C as 50 grams aliquots. To obtain the inclusion bodies, 200 g of bacteria was thawed in 1 liter of lysis buffer supplemented with 2 mg of chicken egg lysozyme per gram of bacteria pellets of was then added and mixed well using a bench-top homogenizer (KitchenAid). The mixture was incubated on ice for 1 hr and sheared with the homogenizer at top-speed for 60 s and cooled in the cold room for 15 min, the shearing process was repeated twice until the solution become less sticky, which was then centrifuged at 10,000 *g*/4°C/15 min. The supernatant was discarded, and the pellets were subjected to a new round of resuspension-shearing-centrifugation process until the pellets became compact and supernatant turned from turbid to translucent. The pellets were finally resuspended in 1 to 2 liters of lysis buffer and aliquot to 20 to 40 50-ml conical tubes, pellet down by centrifugation at 8000 g/10°C/15 min and stored at -20 °C until use.

### I.B. solubilization and refolding screen

To solubilize the I.B., a solubilization buffer containing 50 mM Tris-HCl (pH 8.0), 8 M urea or 6 M guanidine-HCl and 5 mM 2-ME was used. The choice of the solubilization buffer was based on the solubilization effect and contaminating protein level. Taken αBtx for example, this toxin refolded poorly in the presence of contaminating proteins and its I.B. was solubilized well with a solubilization buffer containing 8 M urea. So, after solubilization with 50 mM Tris base, 8 M urea, and 5 mM 2-ME, and centrifuged for 28,000 *g*/10 min/4°C to get rid of insoluble bacteria debris, the pH of the supernatant was adjusted to 8.5 with concentrated HCl solution and further absorbed with Q Sepharose FF media (2 ml of solution/ ml of Q media) equilibrated with 50 mM Tris-HCl (pH 8.5), 8 M urea. Attention should be paid to avoid using high concentrations of reducing chemical reagents (like 100 mM of 2-ME) at the solubilization stage, which will lead to low refolding efficiency that may be caused by blocking the formation of disulfide bonds. After absorption, the I.B. was ready for refolding. For other toxins with higher expression level and more compact inclusion bodies, a solubilization buffer containing 50 mM Tris-HCl (pH 9.0), 6 M guanidine-HCl, 5 mM 2-ME was used.

Refolding condition was optimized with a screening protocol scouting for NaCl concentration (0 or 200 mM), l-cysteine concentration (0-16 mM), l-arginine (0 or 0.5 M), and detergent, such as NDSB-201(0 or 0.2 M), etc. Standard refolding trial was made by diluting 200 μl of I.B. solution into 10 ml of refolding screen solution with detailed recipe presented in Supplementary Figure 1. After refolding, the solutions were left at 4 °C overnight (more than 24 hr), and concentrated each with Amicon Ultra-15 (Millipore, 3 kDa NMWL) ultrafiltration devices to less than 200 μl. The retention was centrifuged at 18,000 g/4°C for 15 min and the supernatant analyzed with non-reducing SDS-PAGE. Generally, mis-paired disulfide bonds could lead to formation of intermolecular disulfide bonds and multimeric species, which are shown as a ladder pattern on non-reducing SDS-PAGE, while correctly paired disulfide bonds facilitate formation of monomeric species, which are usually shown as the smallest band on non-reducing SDS-PAGE. By comparing the yield of monomeric species from different refolding conditions on non-reducing SDS-PAGE, the best refolding condition was selected. The rest of the concentrated solutions were each divided into three parts and dialyzed against low ionic strength buffer with various pH values, such as 20 mM sodium acetate (NaAc, pH 5.0), 20 mM HEPES (pH 7.0, adjusted with NaOH), or 20 mM Tris-HCl (pH 8.0), using a set of home-made micro-dialysis devices (Fiala et al., 2011). Finally, the dialyzed solution was centrifuged at 18,000 *g*/4°C for 15 min. The supernatant was analyzed with non-reducing SDS-PAGE to assess the stability of different refolding species (i.e., monomer and multimers).

### Preparative refolding of rTFPs

After the initial screen, an optimized refolding condition was usually determined, based on the yield of monomeric species seen on non-reducing SDS-PAGE gel. For preparative refolding, freshly solubilized I.B. was poured all in once at a volume ratio of 1:50 into a freshly prepared, ice-cold refolding solution which was stirred rapidly by a magnetic bar throughout the whole process. The refolded solution was left static over one day at 4 °C and concentrated with a compressed nitrogen-gas (or air) driven ultrafiltration device (350 ml Amicon Stirred Cell, 3 kDa NMWL membrane, Millipore), during which time cysteine in the solution gradually react to form cystine and precipitated out and could usually clog the ultrafiltration membrane at the end phase of ultrafiltration, making the process longer. Typically, the refolded solution was either concentrated to a very small volume of several ml (‘not to dry’), or ‘to dry’, leaving no visible liquid on the ultrafiltration membrane, depending on the type of the rTFPs being refolded. For some toxins, like recombinant MTα (rec-MTα), Hannalgesin (rec-Hannagesin), mouse Pate B (rec-mPateB), κ-Bungarotoxin (rec-κBtx), α-Bungarotoxin (rec-αBtx), it is better to concentrate to ‘dry’, which dramatically increased the purity and quality of the final product. For other toxins we tried, such as recombinant mambalgin-1 (rec-Mambalgin-1), mouse and human

Slurp1 (rec-mSlurp1 and rec-hSlurp1), concentrating to dry significantly lowered the final yield. So, trial experiments should be done at this point. To simplify the ultrafiltration process, we designed a ‘storage and oxidation tank and connected in serial in between the compressed gas tank and the stirred ultrafiltration cell (**Figure 1**). The ‘storage and oxidation tank’ not only eliminated the needs for repeated refilling of the stir cell during the ultrafiltration, but was also used as a reaction vessel, guaranteeing complete and homogenous oxidation formation of disulfide bonds under higher dissolved oxygen level under the high pressure. The concentrated product was then re-solubilized with a low ionic strength buffer, which was pre-determined in the dialysis experiment. Normally, proteins with isoelectric point (pI) over 7 was re-solubilized in 30 to 50 ml of 20 mM NaAc (pH 5.0), while proteins with pI less than 7 was solubilized in 20 mM Tris-HCl (pH 8.0) or 20 mM HEPES (pH 7). The solution was then filtered with a 0.2 μm filter and applied to mono S 5 50 GL or mono Q 5 50 GL column, based on the isoelectric point (pI) of the proteins. Bound proteins were eluted with a linear gradient of NaCl to 1 M. The eluted peaks were again analyzed by non-reducing SDS-PAGE. Those eluted later usually contained contaminating proteins, or species inter-connected by intermolecular disulfide bonds. For those not concentrated to ‘dry’ but to small volume, an additional dialysis step was usually added, in which the concentrated solution was dialyzed against the low-ionic strength buffer before being subjected to cation exchange chromatography. For the proteins we tried, a single, large peak was usually seen using the mono S column, and several large peaks were seen using the mono Q column, in which the target species was usually contained in the first peak. At this stage, the refolded rTFP was fairly pure, but for crystal growth, gel filtration was usually done with a Superdex 75 10 300 GL column (GE Healthcare), to further increase the purity of the product and to buffer-exchange to 200 mM ammonium acetate (pH 7°C).

### Native gel shift assay

5 μg of HAP peptide (Harel et al., 2001; Kudryavtsev et al., 2020) were mixed with 5 μg of each of the rTFPs, respectively (with molar ratio HAP:rTFP > 6), incubated at room temperature for 15 min, and run on a 15% native PAGE gel with 50 mM NaAc (pH 5.0) at 120 v/60 min/ 4°C. For the binding assay with the nicotinic acetylcholine receptors, 5 μg of recombinant α-Cobratoxin (rec-αCTX), recombinant Hannalgesin (rec-Hannalgesin), or α-Cobratoxin (αCTX) (Sigma-Aldrich, C6903) was mixed with 5 μg of the recombinant extracellular domain of the α1 subunit of muscle type nicotinic acetylcholine receptor (rec-α1ECD)(Dellisanti et al., 2007; Yao et al., 2002) (molar ratio, rTFP (TFP): α1ECD = 3), incubated on ice for 15 min and run on 12% native gel (standard discontinuous PAGE gel without SDS, 6% for top layer and 10% for bottom layer) with Tris-Glycine buffer (pH 8.3, without SDS) as the running buffer, at 120 v/90 min/4°C. Gels were stained with Coomassie Brilliant Blue G-250 as described(Wittig and Schägger, 2005).

### Labeling of rec-mPate B with fluorescence dye and visualization of binding of rec-mPate B to the mouse spermatozoa

rec-mPate B was labeled with NHS-rhodamine (Thermo Scientific) according to the manufacturer’s recommended protocol. Briefly, 25 μl of rec-mPate B solubilized in PBS (pH 7.4) at 27.2 mg/ml was mixed with 20 mM HEPES (pH 7), 4.13 μl of 18.9 mM NHS-Rhodamine DMSO solution and incubated at room temperature for 60 min, and dialyzed exhaustively against 20 mM HEPES, 0.15 M NaCl. Mouse spermatozoa was obtained as described (Duselis and Vrana, 2007), and was mixed with 1:1000 dilution of the Rhodamine labeled rec-mPate B, washed three times with PBS, and observed under a laser confocal fluorescence microscope.

### X-ray protein crystal diffraction structural validation of rTFPs

For crystallization of rec-αBtx, rec-αBtx was mixed with HAP peptide (Harel et al., 2001) at a molar ratio of 1:1.5, incubated at room temperature for 30 min and then diluted 100 fold with 20 mM NaAc, pH 5.0 and applied to mono S column. Bond protein was eluted with linear gradient of NaCl to 1 M and the sharp peak containing the rec-αBtx-HAP complex was collected, pooled, and concentrated to about 13 mg/ml, dialyzed against 0.1 M HEPES (pH 7.0) exhaustively at 4 °C. For other rTFPs, purified toxin proteins were concentrated to 80 to 150 mg/ml with Amicon Ultra-15 and Amicon Ultra-0.5 (3 kDa NMWL) tubes. Sitting drop crystal screening was done using a robotic system (Crystal Gryphon, Art Robbins Instrument). Hanging drop method was then done manually to optimize the growth condition, by mixing equal volume of well solution and the toxin protein and incubating both at 4 °C and 18 °C. Crystals were then harvested and stored in cryo-conditions and X-ray diffraction data of rec-kBtx, rec-mambalgin 1 and rec-αBtx-HAP complex were collected either with a Rigaku MicroMaxTM-007 home X-ray source coupled with an R-AXIS IV++ image plate. For rec-MTα crystals, X-ray diffraction data was collected at Advanced Photon Source (Argonne National Laboratory, Lemont, IL). The X-ray diffraction data of rec-Hannalgesin and rec-αCTX were collected at Advanced Light Source (Lawrence Berkeley National Laboratory, Berkeley, CA). Data was processed with HKL2000 (Otwinowski and Minor, 1997) or IMosflm (Battye et al., 2011), Pointless, Aimless (Evans and Murshudov, 2013), Ctruncate from the CCP4 suite (Winn

et al., 2011), Molecular Replacement, structure build and refinement was done in Phenix (Liebschner et al., 2019) and Coot (Liebschner et al., 2019). Structural visualization, alignment, and calculation were done in open source PYMOL (Version 2.0 Schrödinger, LLC).

## Results

### Our pipeline is applicable to a wide variety of TFPs with high yield, with 6 rTFPs being structurally validated

Our idea is to use *E. coli* as the workhorse to produce high quality TFPs of biomedical interests. Our pipeline includes codon optimization of the encoding DNA sequence for *E. coli* expression, recombinant protein expression in *E. coli*, isolation of I.B., refolding condition screening, preparative oxidation refolding and purification, structural validation with x-ray diffraction and biochemical methods (**Figure 1**). From construction of the expression vector with known protein or encoding DNA sequences, production of a purified rTFP usually took 4∼5 weeks. For each of the rTFPs, non-reducing SDS-PAGE were carried out to quantify monomeric and multimeric species with inter-molecular disulfide bonds and to check the purity of the final product (**Figure 2**). For a couple of TFPs of various origin (**Supplementary Table 1**), our pipeline was shown to be robust and successful (**Figure 2, Supplementary Material**).

**Figure 2.**
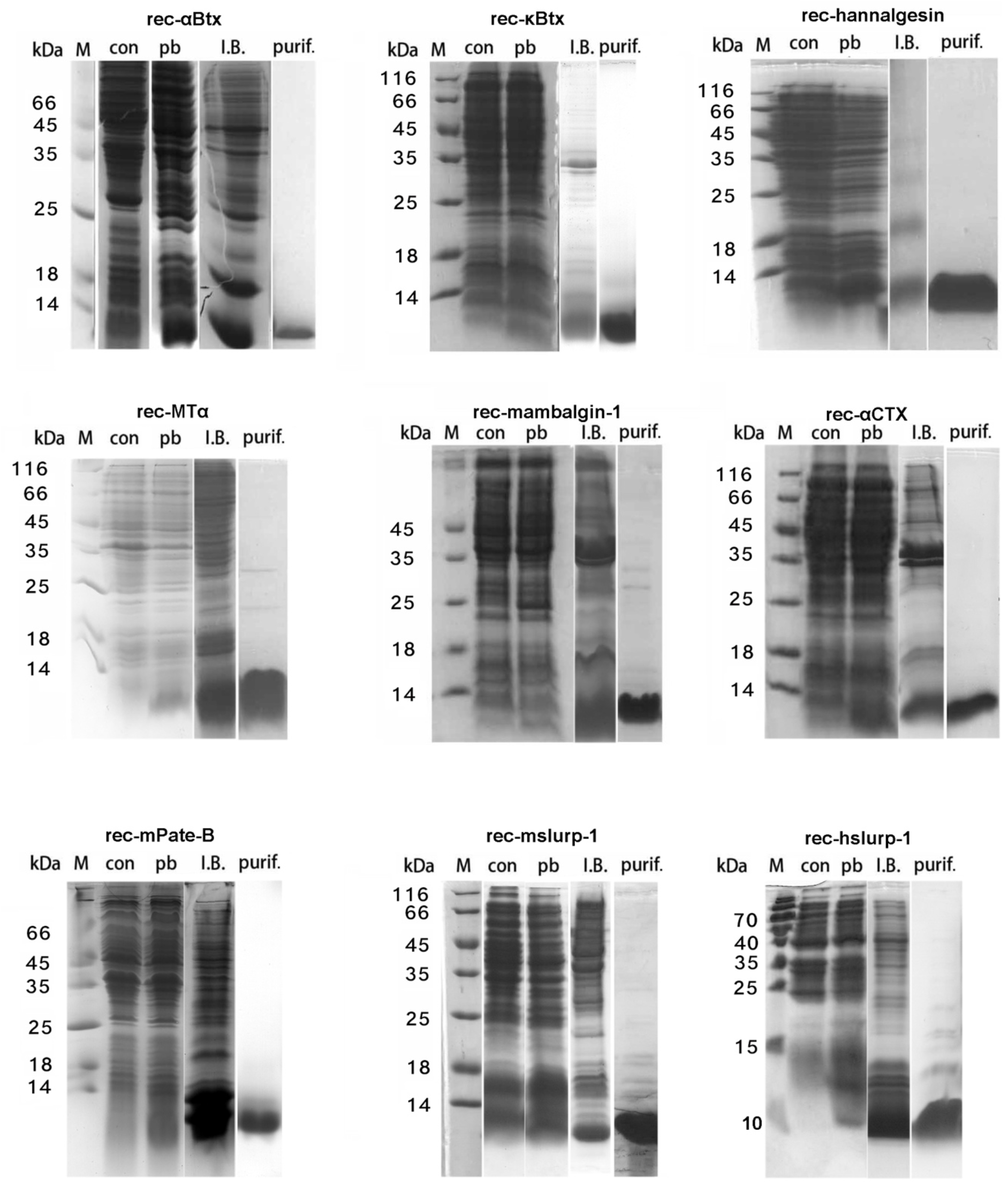
SDS-PAGE analysis of rTFPs at different stages of production. con: control (not induced E. coli cells); pb: IPTG induced E. coli cells; I.B.: isolated inclusion bodies; purif.: purified final product (in non-reducing SDS-PAGE).

To structurally validate the rTFPs, we screened for crystal growth for each rTFP we purified (**Supplementary Material**). Most of our rTFPs’ crystals were formed at very high protein concentrations (**supplementary material**). They were beautiful looking under the microscope with polarized light (**Supplementary Figure 1**), and diffracted x-ray quite well. Six rTFPs’ structures were solved with x-ray crystal diffraction data using molecular replacement with known homologous structures. The statistics for the data were summarized in **Table 1**. From the structural alignment of the solved structures with their native counterparts (such as rec-aBtx-HAP complex, rec-αCTX, rec-kBtx, rec-mambalgin) or with their most homologous native counterparts (such as rec-Hannalgesin and rec-MTα, whose crystal structures were not reported, known homologous TFPs with known structure, such as αCTX and MT1, respectively, were used as the alignment counterpart). Our rTFPs are shown to be almost identical to their natural counterparts, with one or several additional amino acids at the N-terminal, which is a unique mark for their recombinant origin) (**Figure 3**).

**Table 1.** Data collection and refinement statistics.

|  |  | rec-Mambalgin-1 | rec-kBtx | rec-αBtx-HAP | rec-Hannalgesin | rec-MTα | rec-αCTX |
| --- | --- | --- | --- | --- | --- | --- | --- |
| Resolution range (Å) |  | 24.09 - 2.49<br>(2.579 - 2.49) | 36.71 - 1.8<br>(1.864 - 1.8) | 33.35 - 2.4<br>(2.486 - 2.4) | 39.46 - 2.2<br>(2.279 - 2.2) | 29.28 - 1.8<br>(1.865 - 1.8) | 30.77 - 1.57<br>(1.626 - 1.57) |
| Space group |  | P 1 2 1 1 | P 1 | P 1 2 1 1 | P 32 | P 63 | P 3 2 1 |
| Unit cell | Length (Å) | 30.984 80.753 54.032 | 27.0899 30.9899 39.43 | 42.5295 73.1098 79.8397 | 65.5095 65.5095 164.752 | 58.555 58.555 35.395 | 73.9695 73.9695 110.841 |
|  | Angle (°) | 90 90.47 90 | 70.44 75.97 71.94 | 90 103.68 90 | 90 90 120 | 90 90 120 | 90 90 120 |
| Multiplicity |  | 3.9 (4.0) | 4.0 (4.0) | 7.6 (7.7) | 6.1 (6.0) | 9.3(7.2) | 21.7 (20.6) |
| Completeness (%) |  | 94.97 (93.12) | 93.40 (89.76) | 96.32 (94.14) | 99.76 (99.58) | 99.88 (99.84) | 99.83 (99.79) |
| Mean I/sigma(I) |  | 20.12 (7.47) | 16.92 (8.91) | 10.58 (4.53) | 11.67 (3.38) | 53.4 (17.65) | 21.76 (3.96) |
| Wilson B-factor |  | 55.19 | 13.48 | 18 | 41.94 | 21.26 | 20.76 |
| R-merge |  | 0.04624 (0.1469) | 0.05006 (0.1079) | 0.1477 (0.4164) | 0.08822 (0.5762) | 0.049(0.129) | 0.08866 (0.8743) |
| R-meas |  | 0.05376 (0.1683) | 0.05787 (0.1247) | 0.1584 (0.4462) | 0.09657 (0.631) | 0.052(0.137) | 0.09088 (0.8962) |
| CC1/2 |  | 0.997 (0.978) | 0.998 (0.985) | 0.992 (0.943) | 0.996 (0.904) |  | 0.999 (0.951) |
| R-work |  | 0.2179(0.3432) | 0.1658 (0.2067) | 0.2393 (0.2844) | 0.2115 (0.2708) | 0.1897 (0.3007) | 0.1736 (0.2363) |
| R-free |  | 0.2670 (0.3537) | 0.1912 (0.2233) | 0.2750 (0.3626) | 0.2431 (0.3349) | 0.2202 (0.3696) | 0.1844 (0.2515) |
| macromolecules |  | 1845 | 1018 | 2671 | 6094 | 532 | 2040 |
| ligands |  | 15 | 0 | 96 | 25 | 28 | 39 |
| RMS (bonds) |  | 0.008 | 0.007 | 0.006 | 0.007 | 0.009 | 0.012 |
| RMS (angles) |  | 1.18 | 1.01 | 1.09 | 0.98 | 1 | 1.25 |
| Average B-factor |  | 72.89 | 20.55 | 23.99 | 63.69 | 28.06 | 26.68 |
| macromolecules |  | 73.10 | 19.43 | 23.52 | 63.95 | 26.95 | 24.44 |
| ligands |  | 68.76 |  | 63.94 | 72.89 | 47.62 | 51.78 |
| (Values in parentheses are for the highest-resolution shell) |  |  |  |  |  |  |  |

(Values in parentheses are for the highest-resolution shell)

**Figure 3.**
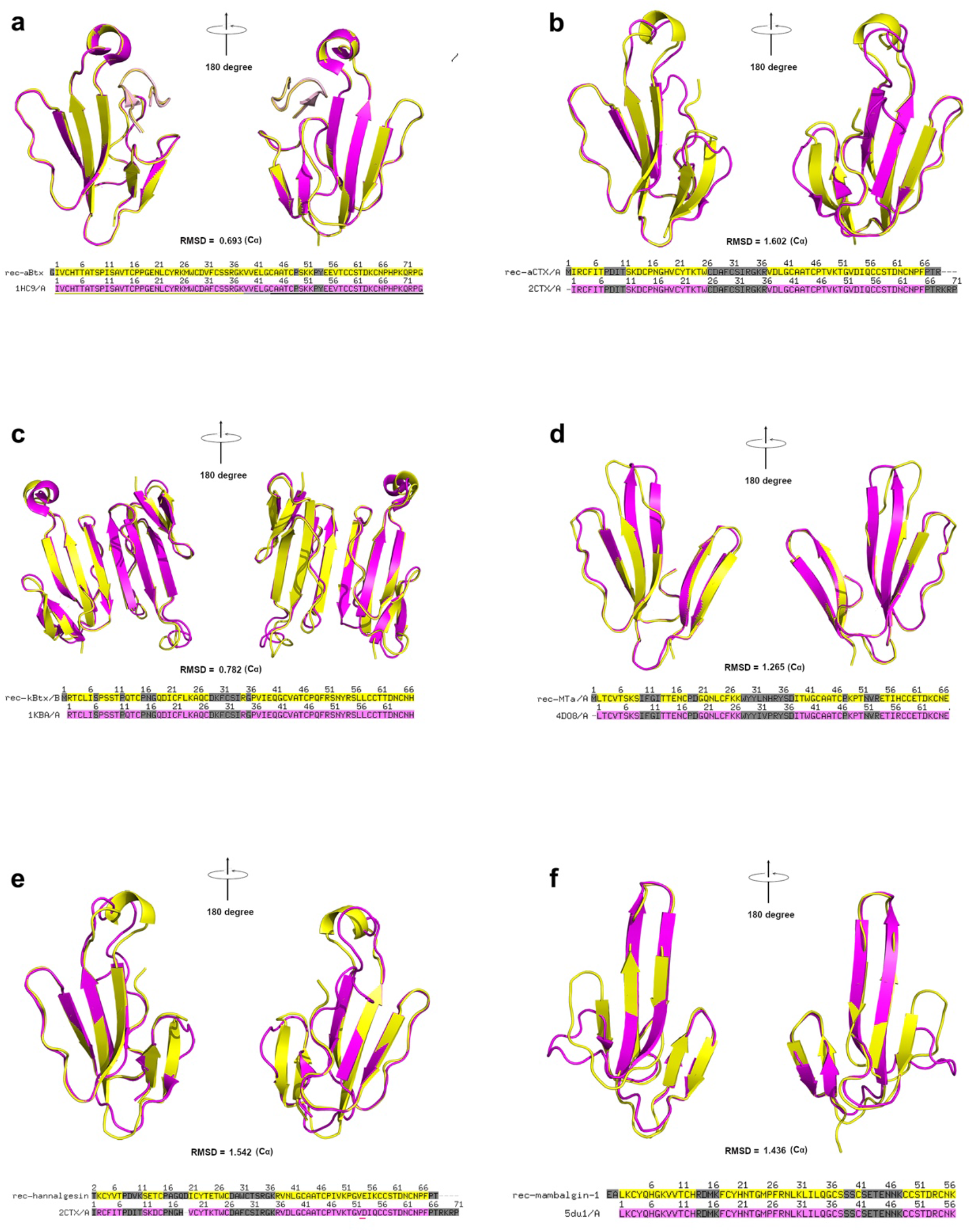
Structural alignment of the crystal structure of rTFPs and their natural counterpart or most homologous natural counterpart. Yellow: rTFP, Magenta: Reported native (homologous) or synthetic counterpart. **a**. rec-αBtx-HAP vs αBtx-HAP; **b**. rec-αCTX vs αCTX; **c**. rec-κBtx vs κBtx; **d**. rec-MTα vs MT1;**e**. rec-Hannalgesin vs αCTX; **f**. rec-Mambalgin-1 vs Mambalgin-1. RMSD was calculated based on the coordinates of Cα of the aligned structures.

Though it was generally considered very difficult to obtain rTFPs using either *E. coli* or other expression systems, with our pipeline, we could repeatedly obtain over one hundred milligrams of rec-MTα, rec-Hannalgesin, rec-αCTX and rec-mPate B, tens of milligrams of rec-mambalgin-1, Slurp1 and milligrams of rec-kBtx and rec-αBtx (**Supplementary material**) through only one round of experiment (usually finished within 4∼5 weeks).

### Most useful scouting conditions for refolding rTFPs are cysteine, salt concentration, and pH

The optimized refolding condition for each recombinant neurotoxin was summarized in the supplementary material. The most critical factors are the concentration of sodium chloride and l-cysteine, and pH value. A weak basic solution, with different NaCl and l-cysteine concentration provided the essential refolding solution. Without Tris base, increasing l-cysteine concentration significantly alter the pH from neutral to acidic, leading to lowered yield of monomeric species (data not shown). L-arginine (Arakawa et al., 2007; Chen et al., 2008; Tischer et al., 2010; Tsumoto et al., 2004) and NDSB-201 (Luca et al., 2012; Wangkanont et al., 2015), two known supplements which are widely used in inclusion body refolding, even though significantly increased the yield of monomeric species in the screening experiment as reflected by non-reducing SDS-PAGE (**Supplementary Figure 2**), lead to formation of a lot of precipitates in the subsequent dialysis removal of these supplements (data not shown), and thus didn’t help much. What’s more, l-arginine and NDSB-201 are relatively expensive and not cost-effective in large scale production. Taken together, L-arginine and NDSB-201 are generally not recommended for refolding rTFPs, at least for those we tried. Normally, rTFPs with high isoelectric point (pI) remained soluble upon challenge with weak acidic solution (such as 20 mM NaAc (pH 5.0)), while certain mammalian three finger toxin-like protein, such as Slurp1, remained soluble only in neutral and slight basic solutions, such as 20 mM HEPES (pH 7.0) and 20 mM Tris-HCl (pH 8.0).

### Complete oxidation is the key to the production of high quality rTFPs

It is common to see I.B. refolding protocols in which people dissolve the I.B. with solutions containing high concentration of reducing agents (such as 100 mM β-mercaptoethanol or 2-ME). While these agents are useful in keeping the free cysteine residue in reduced form and it might not be a problem in certain cases, we found 100 mM 2-ME in I.B. solubilization buffers inevitably lead to failed refolding experiments, which was shown by the extremely low yield and formation of multimeric species (Xu et al., 2015), thus should be avoided when solubilizing the I.B. For correct disulfide bonds pairing between the cysteine residues, a classical and widely used approach is the disulfide shuffling or mixed disulfide bond reactions, in which a predefined redox pairs such as a fixed ratio of reduced-glutathione:oxidized-glutathione, or cysteine:cystine are used (“Disulfide bond formation in proteins,” 1984; Okumura et al., 2011; Qin et al., 2015). In our pipeline, we used a simple, straightforward approach by scouting cysteine and NaCl concentration in screening of refolding conditions, and we noticed that different rTFPs had different sensitivity to cysteine concentration in the refolding experiment (**Supplementary Figure 2)**. We would choose a condition with the lowest concentration of l-cysteine that produced the highest level of monomeric species, as shown by non-reducing SDS-PAGE. In preparative refolding, we used compressed N2 gas and/or air to drive the ultrafiltration device (**Figure 1**). In refolding of rec-kBtx, we found N2 gas was not as good as compressed air, which dramatically decreased the multimeric species in the final product, and only the purified rec-kBtx from this special protocol yielded crystals. Clearly, ultrafiltration with the stirred cell is not only a physical process, but also a biochemical process in which dissolved oxygen level is critical for the correct and complete formation of disulfide bonds. Considering this, we designed a storage tank (**Figure 1**), which acted as a oxidation reservoir for the refolded mixture and gas, eliminate the needs for opening the ultrafiltration cell to refill and also to adjust the ratio between air and N_2_. A typical preparative ultrafiltration procedure took about 4 weeks, during the last a few days of which a large amount of white precipitate (which turned out to be cystine, the oxidized form of cysteine) showed up in the concentrated solution, which were found to be a good sign of complete oxidation, since most of our high quality rTFP were produced in this way. Some of our rTFPs were tested for their stability by prolonged storage at 4 °C, and were shown to be stable even after one year of storage at 4 °C, and only trace amounts of dimeric and multimeric species were found (**Figure 4a**), consistent to previous report about the stability of natural TFNs (Nirthanan et al., 2015), and prove the high quality of the rTFP from our pipeline.

**Figure 4.**
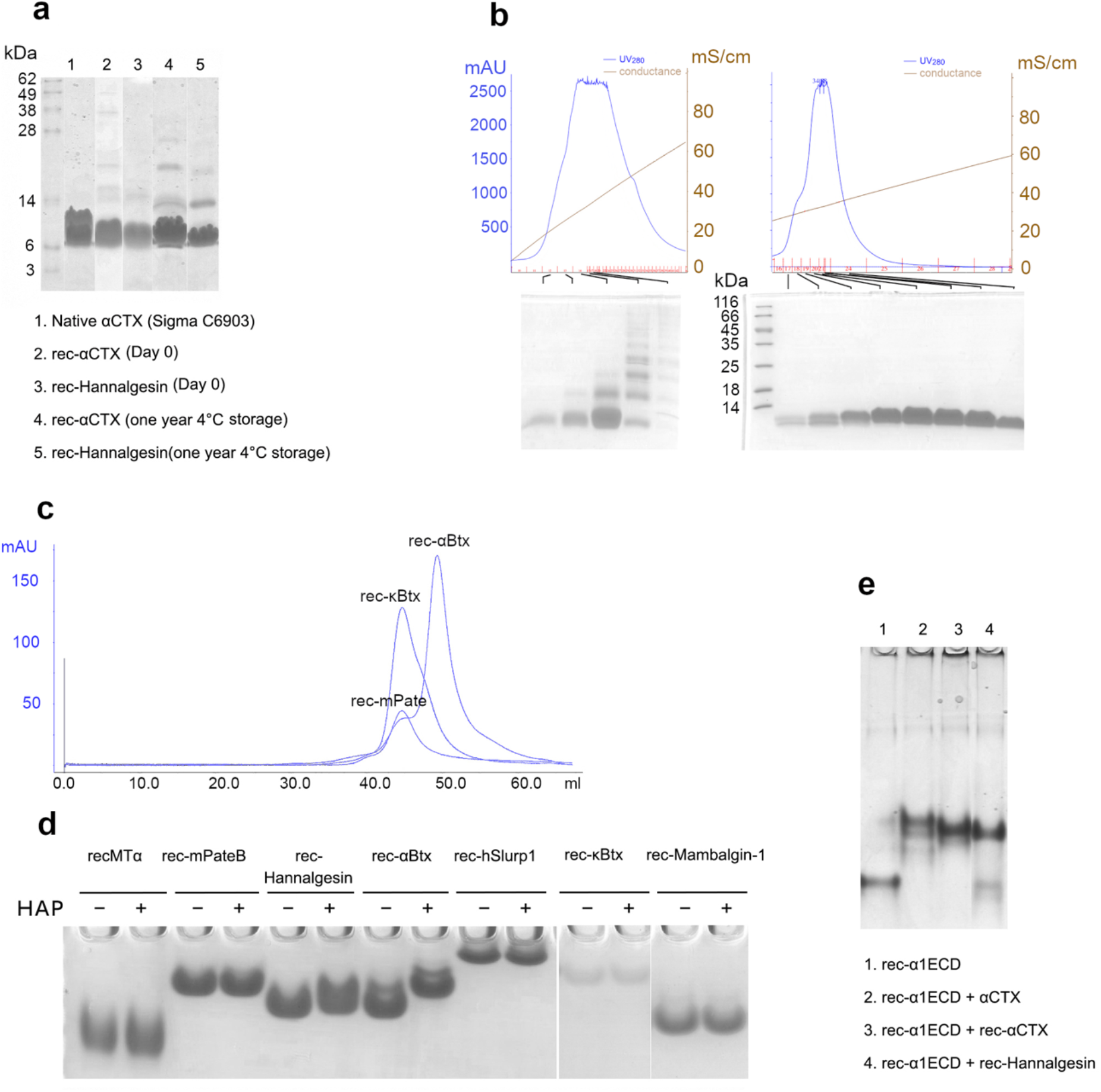
Biochemical characterization of the rTFPs. **a**. Stability of rec-αCTX and rec-Hannalgesin upon prolonged storage at 4 °C, analyzed with non-reducing SDS-PAGE; **b**. ‘Concentrate to dry’ strategy (right) efficiently removed multimeric species that were hard to separate with mono S column (left, in which the refolded product was not ‘concentrate to dry’; **c**. Gel filtration analysis of rec-αBtx, rec-mPate B and rec-κBtx. **d**. Native gel shift assay of various rTFPs with HAP peptide. **e**. Native gel shift assay of rec-α1ECD with native αCTX, rec-αCTX and rec-Hannalgesin.

### Concentrate to dry is a simple and efficient step for removing incorrectly folded species

It is interesting to note this point, since we found that multimeric species, which were generally regarded as incorrectly folded product with wrong pairing of disulfide bonds, were inevitable as a part of refolding product and hard to be completely separated from the correctly folded species using common chromatography approaches, such as gel filtration (data not shown) and ion exchange. However, it turned out that ultrafiltration of the refolded product to dry dramatically increased the purity for some rTFPs (**Figure 4b**). It is possible that incorrectly folded but soluble rTFPs tend to form insoluble aggregates only when they are concentrated to extremely high concentrations, which was achieved in the “concentrate to dry” approach. It is thus noteworthy to try two ultrafiltration strategies, “to dry, or not to dry”, which in most cases could make a great difference.

### Recombinant rTFPs shows good, expected elution profile in gel filtration chromatography and were shown to be active in biochemical and morphological assays

We compared the elution volume of rec-αBtx, rec-κBtx, rec-mPate B in gel filtration column (Superdex 75 10 300 GL, GE Healthcare), and found that rec-αBtx, which were known as a monomer, eluted much later than rec-κBtx, which were known as a dimmer, while rec-mPate B elute at similar volume as rec-κBtx, suggesting rec-mPate B also form a dimmer (**Figure 4c**), which is in accordance with earlier reports (Dewan et al., 1994) that κBtx exists in dimeric form and also with the solved crystal structures (**Figure 3c**). To test the binding specificities of the rTFPs, HAP peptide, a known peptide derived from the nicotinic acetylcholine receptor (Harel et al., 2001), was mixed with various rTFPs and separated on a 15% native PAGE gel at pH 5.0. HAP peptide was only able to shift rec-αBtx and only slightly shift rec-Hannalgesin, but not rec-MTα, rec-mPate B, rec-κBtx, rec-Mambalgin-1 and rec-hSlurp1 (**Figure 4d**). Also, rec-αCTX and rec-Hannalgesin was shown to bind the extracellular domain of α1 subunit of the nicotinic acetylcholine receptor (rec-α1ECD) (Dellisanti et al., 2007), like the native αCTX isolated from Naja Kaouthia (**Figure 4e**).

To test the binding activity of rec-mPate B to sperm, we labeled rec-mPate B with NHS-rhodamine and visualized the binding of rec-mPate B to spermatozoa freshly isolated from the epididymis of the mouse under the fluorescence microscope. The preliminary result suggested binding of rec-mPate B to the head and tail of mouse spermatozoa (**Supplementary Figure 3)**

## Discussion

TFPs are a large collection of proteins with important functions and applications. Traditionally, such proteins were isolated from the venom of the snakes, with very few recombinantly obtained in the lab with in-depth analysis and verification. Because of their scarcity, unique properties and applications, these proteins are very expensive (at the level of hundreds to thousands of US dollars per milligrams) and some are not commercially available. κBtx, for example, a unique α_3_β_2_ nicotinic acetylcholine receptor binder, is not commercially available (personal communications). Because TFPs usually contain 4 to 5 pairs of disulfide bonds, it is usually very hard to recombinantly express them, and those commercially available are mostly purified from snake venoms. Some researchers used chemical synthesis that successfully obtained these rTFPs, such as mambalgin-1 and mambalgin-2 (Diochot et al., 2012; Mourier et al., 2016; Pan et al., 2014; Salinas et al., 2021; Schroeder et al., 2014; Sun et al., 2018). However, due to the high cost in chemical synthesis and limited yields, these successful attempts did not change the overall scenario for production of other TFPs.

With our pipeline, however, milligrams to hundreds of milligrams of rTFPs could be obtained in the lab. Through extensive biochemical assays and structural analysis, we were able to show our rTFPs were almost identical to their native counterparts. Since several of our rTFPs reached milligrams to hundreds of milligrams on a single lab-scale production cycle, these rTFP could thus replace their natural counterparts, and the pipeline is worth to be exploited for production of other TFPs further, which is of general interest in the field.

## Contributions

J.X. conceived the idea and designed the experiments. J.X. and J.L. constructed the expression plasmids, J.X. did the recombinant expression, refolding, and purification experiments. J.X., A.L. did the crystallization screen. J.X. harvested the crystals and collected the diffraction data. J.X., X.L, S.L. and L.C. solved the structure and did the structural analysis.

## Data availability

Coordinates and structure factors have been deposited to the Protein Data Bank with accession number of 7ULB (rec-Mambalgin-1), 7ULR (rec-kBtx), 7ULP (rec-αBtx-HAP complex), 7ULQ (rec-Hannalgesin), 7ULS (rec-MTα) and 7ULG (rec-αCTX).

## Conflict of Interest

There are no competing interests between the authors.

## Acknowledgement

This research used resources of the Advance Light Source, which is a DOE Office of Science User Facility under contract no. DE-AC02-05CH11231. This research used resources of the Advanced Photon Source, which is a DOE office of Science User Facility under contract no. DE-AC02-06CH11357. We thank Advanced Light Source beamline 8.2.1 and Advanced Photon Source beamline 23ID-D and 19ID-D staff members and scientists for help with data collection. We thank Dr. Aaron Wolfe for help with data collections. The research is supported in part by National Institutes of Health grants R01GM064642.

---

**Supplementary Table 1.** Properties of various TFPs.

| TFP name | Origin | Specificity | No. of disulfide bonds | Known structure? | Reference |
| --- | --- | --- | --- | --- | --- |
| **αBtx** | Bungarus Multicinctus | $\alpha 1\beta 1\gamma(\epsilon)\delta$ nAChR blocker | 5 | Yes | (Chang and Lee, 1963; Harel et al., 2001; Love and Stroud, 1986) |
| **κBtx** | Bungarus Multicinctus | $\alpha 3\beta 2$ nAChR blocker | 5 | Yes | (Cartier et al., 1996; Chiappinelli, 1983; Dewan et al., 1994; Fiordalisi et al., 1994, 1991; Loring and Zigmond, 1988; Luetje et al., 1990) |
| **Hannalgesin** | Ophiophagus hannah | NOS activator<br>$\alpha 2\beta\gamma\epsilon$ nAChR blocker | 5 | No | (Pu et al., 1995a, 1995b) |
| **Mambalgin-1** | Dendroaspis polylepis | ASIC1a blocker | 4 | Yes | (Diochot et al., 2012; Mourier et al., 2016; Salinas et al., 2021, 2014; Sun et al., 2018) |
| **MTa** | Dendrosaspis angusticeps | $\alpha 2B$-adrenoceptor blocker | 4 | No | (Koivula et al., 2010) |
| **αCTX** | Naja kaouthia | $\alpha 2\beta\gamma\epsilon$ nAChR blocker | 5 | Yes | (Betzel et al., 1991; Karlsson and Eaker, 1972) |
| **mouse Pate B** | Mouse | phosphatidylethanolamine phosphatidylserine? | 5 | No | (Levitin et al., 2008; Luo et al., 2001; Turunen et al., 2011) |
| **hSlurp1** | Human | $\alpha 7$ nAChR? | 5 | No | (Grønlien et al., 2007; Throm et al., 2018) |
| **mSlurp1** | Mouse | ? | 5 | No | (Swamynathan et al., 2012; Upadhyay, 2019) |

**Supplementary Figure 1.**
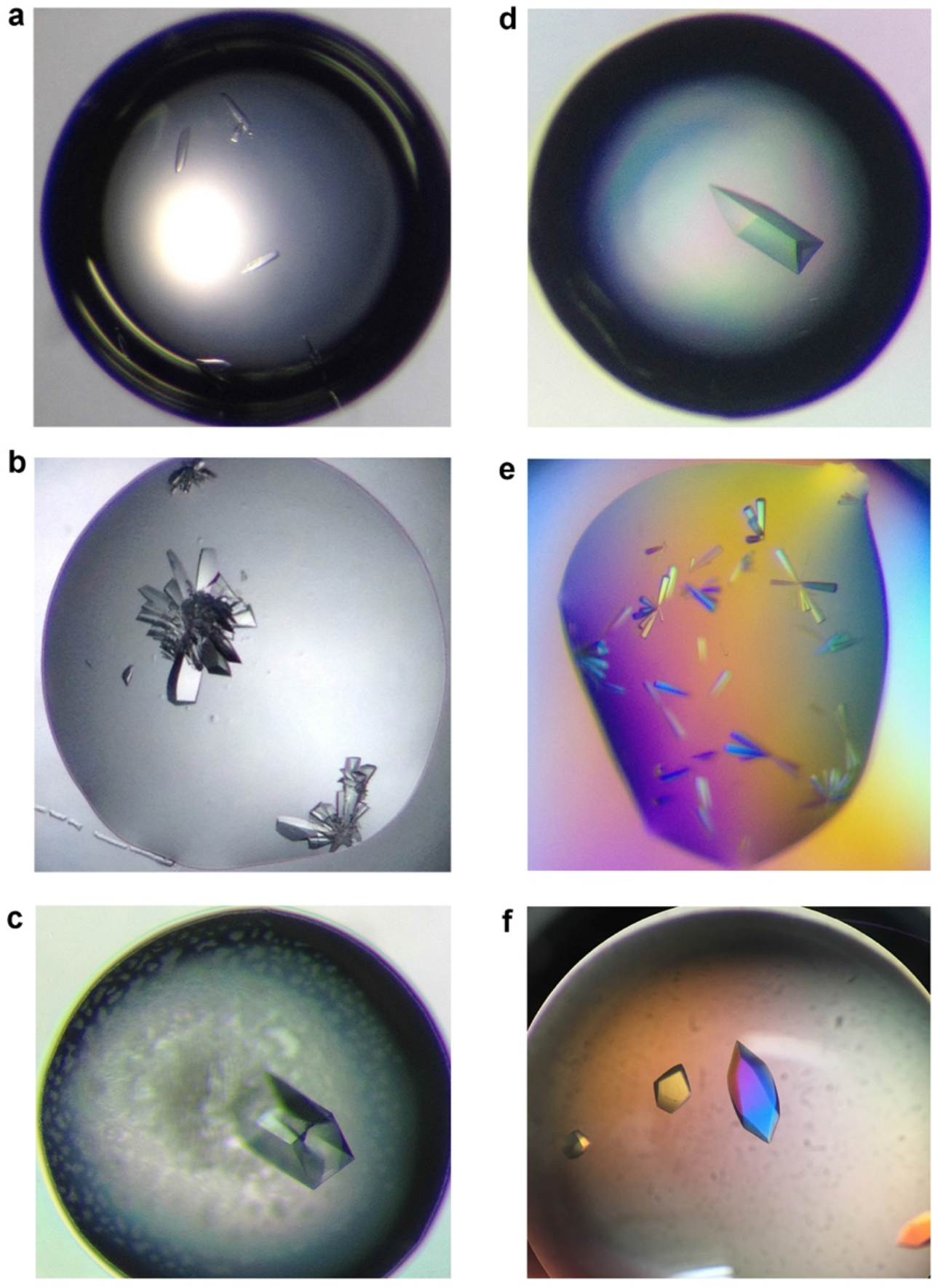
Microscopic view of protein crystals from various rTFPs. **a**. rec-αBtx-HAP complex; **b**. rec-κBtx; **c**. rec-Mambalgin-1; **d**. rec-Hannalgesin; **e**. rec-MTα; **f**. rec-αCTX

**Supplementary Figure 2.**
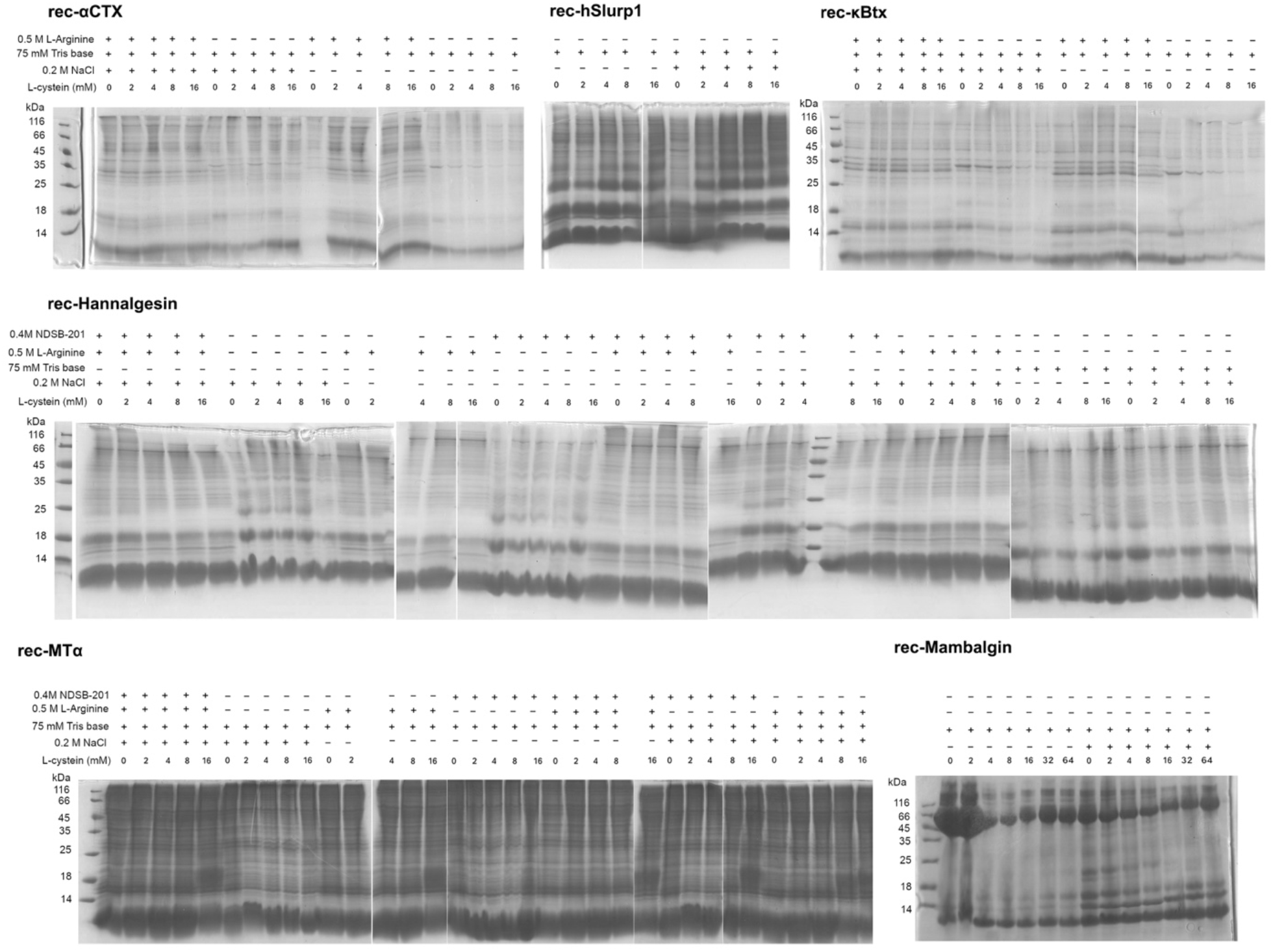
Refolding condition screening of rTNFs Non-reducing SDS-PAGE analysis of refolding products from various conditions were concentrated and analyzed by 15% non-reducing SDS-PAGE.

**Supplementary Figure 3.**
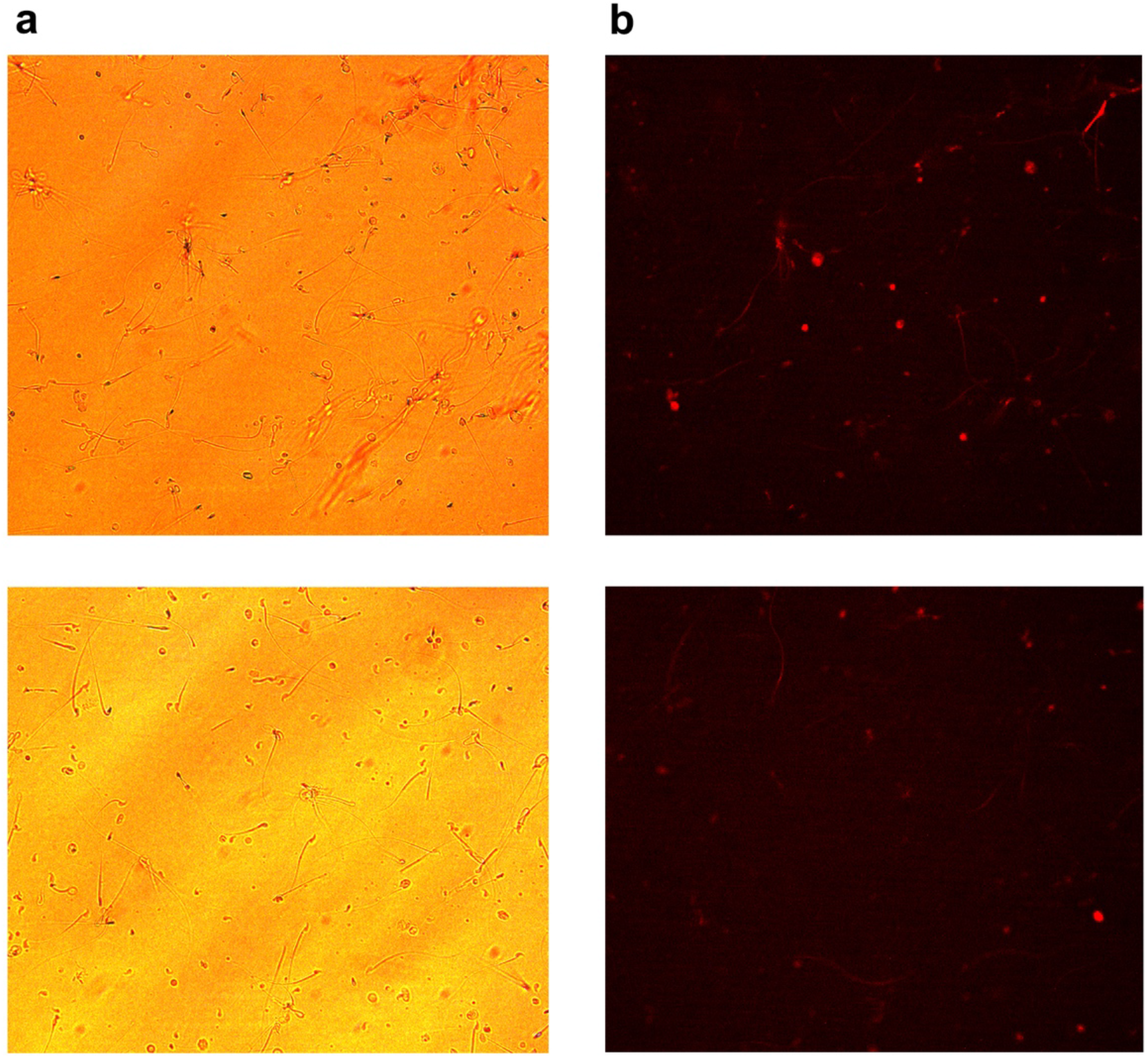
Fluorescence microscopic picture showing the binding of rec-mPate B to the spermatozoa from mouse epididymis. Bright field microscopic picture of mouse spermatozoa (**a**) and fluorescent graph of possible binding of rhodamine-labeled rec-mPate B to the head and tail of the spermatozoa (**b**) under the same view.

## Supplemented Material. Coding DNA sequences, a.a. sequences, protein properties, refolding conditions and key points and crystallization conditions for rTFPs

### rec-αBtx (V31)

#### Coding sequence

atgggtATTGTCTGTCACACTACGGCAACGAGTCCGATCAGCGCAGTTACGTGCCCGCCGGGTGAAAACCTGTGTTATCGTAAAATGTGGTGCGATGTGTTTTGTAGCTCTCGCGGTAAAGTGGTTGAACTGGGT TGCGCAGCAACCTGTCCGAGCAAAAAACCGTACGAAGAAGTTACCTGCTGTTCTACGGATAAATGTAATCCGCATCCGAAACAGCGTCCGGGTTAA

### Translated protein sequence

MGIVCHTTATSPISAVTCPPGENLCYRKMWCDVFCSSRGKVVELGCAATCPSKKPYEEVTCCSTDKCNP HPKQRPG

**Protein parameter:** Number of amino acids: 76; Molecular weight: 8210.54; Theoretical pI: 8.36

**Refolding result:** good

**I.B. solubilization solution**: 75 mM Tris-HCl, pH 8.8, 8 M urea, 10 mM 2-ME

**Refolding solution**: 50 mM Tris-HCl, 16 mM l-cysteine, 0.2 M NaCl.

**Key process**: 1. Purify I.B. with Q Sepharose FF before refolding, and buffer exchanged to 50 mM Tris-HCl, pH 8.8, 8 M urea before refolding. 2. Ultrafiltrate to dry.

**Crystalization condtion:** mono S 5/50GL column purified rec-αBtx-HAP complex, buffer exchanged by dialysis to 0.1 M HEPES(pH 7.5), OD_280_= 11.04. 1:1 with 0.1 M HEPES (pH 7.5), 35% PEG3350, 0.2 M MgCl_2_. 18 °C

**X-ray diffraction data collection:** yes

**Structural solution:** yes

### rec-αCTX

#### Coding sequence

atgATCCGTTGCTTCATCACCCCGGACATCACCTCTAAAGACTGCCCGAATGGCCACGTCTGCTACACGAAAACCTGGTGCGACGCTTTCTGCTCTATCCGTGGTAAACGTGTTGACCTGGGTTGCGCTGCTACC TGCCCGACCGTTAAAACCGGTGTTGACATCCAGTGCTGCTCTACCGACAACTGCAACCCGTTCCCGACCCGTAAACGTCCGTAA

**Translated protein sequence**: MIRCFITPDITSKDCPNGHVCYTKTWCDAFCSIRGKRVDLGCAATCPTVKTGVDIQCCSTDNCNPFPTRK RP

**Protein parameter:** number of amino acids: 72; Molecular weight: 7962.2; Theoretical pI: 8.59

**Refolding result:** good

**I.B. solubilization solution**: 50 mM Tris-HCl, pH 8.8, 6 M Guanidine-HCl, 5 mM 2-ME.

**Refolding solution**: 75 mM Tris base, 8 mM cysteine, 0.2 M NaCl.

**key process**: 1. Ultrafiltrate to dry. 2. Oxidization should be complete for emergence of large amount of white cystine crystalline.

**Yield:** 1 mg/ g bacteria (wet pellet)

**Crystallization condition**: 110 mg/ml in 200 mM NH_4_Ac (pH 7.0/25 °C), 1:1 with 0.1 M HEPES (pH 7.9), 30% Jeffamine M-600 (pH 7.0), 18 °C

**X-ray diffraction data collection:** yes

**Structural solution:** yes

### rec-κBtx

#### coding sequence

atgCGTACCTGTCTGATTAGCCCGTCCAGCACCCCGCAAACCTGTCCGAATGGTCAAGATATTTGTTTTCTGAAGGCCCAGTGTGATAAATTTTGCAGCATTCGTGGCCCGGTGATCGAACAGGGTTGCGTTGC GACCTGTCCGCAATTTCGCTCTAACTACCGTTCACTGCTGTGCTGTACCACCGACAACTGTAATCATT AA

#### Translated protein sequence

MRTCLISPSSTPQTCPNGQDICFLKAQCDKFCSIRGPVIEQGCVATCPQFRSNYRSLLCCTTDNCNH

**Number of amino acids**: 67; Molecular weight: 7406.5; Theoretical pI: 8.07

**Refolding Result:** good

**I.B. solubilization solution**: 50 mM Tris-HCl, pH 8.8, 8 M urea, 10 mM 2-ME

**Refolding solution**: 75 mM Tris Base, 16 mM cysteine, 0.2 M NaCl.

**Key process:**

1. Use compressed air to drive the Amicon Stirred Cell (200 ml).
2. Freezing and thawing the inclusion body solution (in 50 mM Tris-HCl, pH 8.8, 8 M urea, 10 mM 2-ME) and centrifugation to get rid of contaminating protein before refolding.
3. Ultrafiltrate to dry.

**Yield:** 0.1 mg purified product / g bacteria (wet pellet)

**Crystalization condtion:**117 mg/ml in 200 mM NH_4_Ac, pH 7.0, 1:1 with 0.15 M DL-malic acid pH 7.0, 20% PEG 3350 at 4 °C

**X-ray diffraction data collection:** yes

**Structural solution:** yes

### rec-MTα

#### Coding sequence

atgCTGACCTGCGTTACCTCCAAATCTATCTTCGGCATCACGACGGAAAACTGCCCGGACGGCCAGAACCTGTGCTTCAAAAAGTGGTATTATCTGAACCATCGTTACAGCGATATTACGTGGGGTTGCGCAGC AACCTGTCCGAAACCGACGAACGTGCGCGAAACCATCCACTGCTGTGAAACCGACAAGTGCAATGA ATAA

#### Translated protein sequence

MLTCVTSKSIFGITTENCPDGQNLCFKKWYYLNHRYSDITWGCAATCPKPTNVRETIHCCETDKCNE

Number of amino acids: 67 Molecular weight: 7684.7 Theoretical pI: 6.68.

**I.B. solubilization solution**: 50 mM Tris-HCl, pH 8.8, 6 M Guanidine-HCl, 5 mM 2-ME

**Refolding result:** good

**refolding solution**: 75 mM Tris base, 8 mM cysteine, 0.2 M NaCl.

**key process**: No.

**Yield:** 1∼ 2 mg purified product / g bacteria (wet pellet)

**Crystallization condition**: 128 mg/ml in 200 mM NH_4_Ac (pH 7.0), 1:1(v/v) with 1.26 M sodium phosphate monobasic monohydrate, 0.14 M potassium phosphate, pH 5.6 at 18 °C

**X-ray diffraction data collection:** yes

**Structural solution:** yes

### rec-Mambalgin-1

#### coding sequence

atgAAGAGAGAAGCTGAAGCCTTAAAGTGCTATCAACACGGTAAAGTCGTAACCTGCCACAGAGACATGAAGTTCTGCTATCACAACACAGGTATGCCTTTTAGAAATTTGAAGTTGATATTGCAAGGTTGTTCTTC ATCCTGCTCTGAAACTGAAAACAATAAGTGCTGCTCCACCGACAGATGTAACAAAGGTTCA

**Translated protein sequence**: MKREAEALKCYQHGKVVTCHRDMKFCYHNTGMPFRNLKLILQGCSSSCSETENNKCCSTDRCNKGS

**Number of amino acids:** 66; **Molecular weight:** 7522.65; **Theoretical pI:** 8.87

**Refolding result:** good

**I.B. solubilization solution**: 50 mM Tris-HCl, pH 9.0, 6 M Guanidine-HCl, 5 mM 2-ME

**Refolding solution**: 75 mM Tris base, 16 mM cysteine, 0.2 M NaCl

**Yield:** 0.2∼0.5 mg purified product / g bacteria (wet pellet)

**Key process**: **NOT** concentrate to dry (would be hard to resolubilize if concentrate to dry), instead, dialyze the retention against 20 mM sodium acetate, pH 5.0 to remove contaminated proteins and multimeric species, which would precipitate out

**Crystallization condition**: rec-Mambalgin-1(122 mg/ml in 200 mM NH4Ac (pH 7.0)), 1:1 (v/v) with 0.1 M HEPES, pH 7.0, 32% Jeffamine M600, 0.1 M KSCN, 18 °C

**X-ray diffraction data collection:** yes

**Structural solution:** yes

### rec-Hannalgesin

#### Coding sequence

atgACGAAATGCTACGTTACCCCGGATGTTAAAAGCGAAACCTGCCCGGCTGGTCAAGATATTTGCTACACGGAAACCTGGTGCGATGCGTGGTGCACCAGCCGTGGCAAACGCGTCAACCTGGGTTGCGCGG CCACGTGTCCGATTGTGAAACCGGGCGTTGAAATCAAATGCTGCTCCACCGACAACTGTAACCCGTT CCCGACCCGCAAACGCCCGTAA

#### Translated protein sequence

MTKCYVTPDVKSETCPAGQDICYTETWCDAWCTSRGKRVNLGCAATCPIVKPGVEIKCCSTDNCNPFPT RKRP

**Number of amino acids**: 73 Molecular weight: 8050.3 Theoretical pI: 8.36

**Refolding result:** good

**I.B. solubilization solution**: 50 mM Tris-HCl, pH 9.0, 6 M Guanidine-HCl, 5 mM 2-ME

**Refolding solution**: 75 mM Tris base, 16 mM cysteine, 0.2 M NaCl

**Yield:** 1∼2 mg purified product / g bacteria (wet pellet)

**Key process**: Concentrate to dry (would efficiently remove contaminated proteins and multimeric species) and resolubilize the peptide with 20 mM sodium acetate, pH 5.0

**Crystallization condition**: rec-hannalgesin at 80 mg/ml in 200 mM NH_4_Ac (pH 7.0), 1:1 with 0.1 M Bis-Tris, 0.2 M (NH_4_)_2_SO_4_, 25% PEG3350, at 18 °C

**X-ray diffraction data collection:** yes

**Structural solution:** yes

### rec-mSlurp1 (recombinant mouse Slurp1)

#### Coding sequence

atgTTTCGCTGCTATACCTGTGAACAACCGACGGCTATCAACTCATGTAAAAATATCGCTCAATGTAAAATGGAAGACACCGCCTGCAAAACCGTGCTGGAAACGGTTGAAGCGGCCTTTCCGTTCAACCATTCC CCGATGGTCACCCGTAGCTGCAGCTCTAGTTGTCTGGCAACGGATCCGGACGGCATTGGTGTTGCG CACCCGGTGTTCTGCTGTTTCCGTGACCTGTGTAACTCTGGTTTTCCGGGCTTTGTGGCGGGCCTGT AA

#### Translated protein sequence

MFRCYTCEQPTAINSCKNIAQCKMEDTACKTVLETVEAAFPFNHSPMVTRSCSSSCLATDPDGIGVAHPV FCCFRDLCNSGFPGFVAGL

**Number of amino acids:** 89; **Molecular weight:** 9594.04; **Theoretical pI:** 5.47

**Refolding result:** good

**I.B. solubilization solution**: 50 mM Tris base, 8 M urea, 5 mM 2-ME

**Refolding condition**: 50 mM Tris-HCl (pH 9.0), 4 mM cysteine

**Key process**: concentrate NOT to dry, dialyze against 10 mM HEPES, pH 7.5 and purified with mono Q column

**Crystallization condition**: 128 mg/ml with equal 0.8 M Succinic acid pH 7.0 at 4 °C

**X-ray diffraction data collection:** No

### rec-hSlurp1(recombinant human Slurp1)

#### coding sequence

atgCTGAAATGCTACACCTGCAAAGAACCGATGACCTCTGCTTCTTGCCGTACCATCACCCGTTGCAAACCGGAAGACACCGCTTGCATGACCACCCTGGTTACCGTTGAAGCTGAATACCCGTTCAACCAGTCT CCGGTTGTTACCCGTTCTTGCTCTTCTTCTTGCGTTGCTACCGACCCGGACTCTATCGGTGCTGCTC ACCTGATCTTCTGCTGCTTCCGTGACCTGTGCAACTCTGAACTGTAA

#### Translated protein sequence

MLKCYTCKEPMTSASCRTITRCKPEDTACMTTLVTVEAEYPFNQSPVVTRSCSSSCVATDPDSIGAAHLIF CCFRDLCNSEL

**Number of amino acids**: 82; **Molecular weight**: 8984.3; **Theoretical pI**: 5.15

**Refolding Result:** good

**I.B. solubilization solution**: 50 mM Tris base, 8 M urea, 5 mM 2-ME

**Refolding condition**: 50 mM Tris-HCl, 0.2 M NaCl, 4 mM cysteine

**Key process**: 1. Fusion of 6 x Histidine tag would significantly reduce refolding efficiency; 2. **NOT** concentrate to dry after refolding, dialyze against 10 mM HEPES, pH 7.5 and purified with mono Q column

**Crystallization condition**: no crystal obtained

### rec-mPate-B(recombinant mouse Pate B)

#### Coding sequence

atgCTGATCTGCAACTCTTGCGAAAAATCTCGTGACTCTCGTTGCACCATGTCTCAGTCTCGTTGCGTTGCTAAACCGGGTGAATCTTGCTCTACCGTTTCTCACTTCGTTGGTACCAAACACGTTTACTCTAAACA GATGTGCTCTCCGCAGTGCAAAGAAAAACAGCTGAACACCGGTAAAAAACTGATCTACATCATGTTC GGTGAAAAAAACCTGATGAACTTCctcgag<u>CACCACCACCACCACCACTGA</u>

#### Translated protein sequence

MLICNSCEKSRDSRCTMSQSRCVAKPGESCSTVSHFVGTKHVYSKQMCSPQCKEKQLNTGKKLIYIMFG EKNLMNFLEHHHHHH

**Number of amino acids**: 84; **Molecular weight:** 9683.22; **Theoretical pI:** 9.08

**Refolding Result:** good

**I.B. solubilization solution**: 5 mM imidazole, 6 mM Guanidine-HCl, 10 mM 2-ME

**Refolding condition**: 50 mM Tris-HCl, pH 9

**Key process**: 1. Fusion of 6xHistidine tag did not significantly reduce refolding efficiency; 2. Concentrating to dry after refolding would efficiently remove contaminated proteins and multimeric species) and resolubilize the peptide with 20 mM sodium acetate, pH 5.0 and purified with mono S column

**Crystallization condition**: no crystal obtained

## Notes

### Competing Interest Statement

The authors have declared no competing interest.

